# AbAgym: a well-curated dataset for the mutational analysis of antibody-antigen complexes

**DOI:** 10.1101/2025.07.15.664862

**Authors:** Gabriel Cia, Dong Li, Simon Poblete, Marianne Rooman, Fabrizio Pucci

**Author notes:** Contributed Equally to this work (Gabriel Cia, Dong Li).

## Abstract

With monoclonal antibodies becoming one of the largest classes of biopharmaceuticals, it is important to have curated data to train computational models that can accelerate their design. Despite the massive amount of mutagenesis data generated on antibody-antigen interactions, only a few small, well-curated datasets are available. This paper introduces AbAgym, a manually curated dataset comprising approximately 335k mutations in antibody-antigen complexes, including one tenth of interface mutations, whose effects on antibody-antigen binding have been experimentally quantified through deep mutational scanning (DMS) experiments. We collected and curated 67 DMS datasets from the literature together with the three-dimensional structure of each antibody-antigen complex. We benchmarked the performance of established force field methods as well as recent machine learning models that predict the change in binding affinity upon mutation. The former achieved modest performance, whereas the latter performed only marginally better than random. Finally, our analysis of hotspot residues responsible for immune evasion highlights the importance of accounting for biological complexities, such as conformational changes or oligomeric states that influence antibody-antigen binding, which are often overlooked. Abagym is freely available for academic use at https://github.com/3BioCompBio/Abagym.

## Introduction

Antibodies play a central role in the human immune system by binding to immunogenic molecules with high affinity and specificity, making them versatile tools widely used in the fields of, e.g., biomedical diagnostics and drug development [1, 2]. The binding mechanism of an antibody to immunogenic molecules, particularly protein antigens, is primarily driven by residues within the antibody’s complementarity-determining regions (CDRs) that interact with key residues on the antigen’s surface, commonly referred to as hotspot residues [3, 4].

The determination of how single-site mutations affect antibody-antigen binding affinity has become an important area of research. It not only drives advances in the rational design and optimization of monoclonal antibodies [1, 2], but also provides critical insights into how variants escape antibody-mediated immune responses in viral infections [5, 6], bacterial infections [7] and cancer [8, 9]. A series of experimental methods have been developed for this task, among which DMS experiments [10, 11]. A large number of DMS studies have been published to date, particularly during the COVID-19 pandemic, typically aiming to identify antigen residues that are the most crucial for antibody binding and therefore represent potential sites for immune escape mutations [12, 13, 14, 15].

Despite the vast amount of antibody-antigen DMS data generated so far, there is no comprehensive repository that has collected, curated, and integrated these datasets. To fill this gap, we developed AbAgym, a large-scale dataset with data from 67 antibody-antigen DMS experiments, which we manually curated from their respective scientific publications. The ensemble of datasets cover different types of phenotypic scores and various antigen types, including the SARS-CoV-2 spike protein, the human immunodeficiency virus (HIV) envelope protein, lysozyme, nerve growth factors (NGF), etc. Importantly, we also paired each DMS dataset with the 3-dimensional (3D) structure of the antibody-antigen complex by using experimentally determined structures from the Protein Data Bank (PDB) [16] when available, and homology models [17] when, as is most often the case, the exact DMS sequence cannot be directly mapped to any structure available in the PDB.

Thanks to the massive amount of mutagenesis data collected, AbAgym can play a pivotal role in advancing our understanding of antibody-antigen interfaces and in identifying key residues and interactions that are crucial for protein-protein complex formation. Owing to its careful manual curation, AbAgym can be used by the community to train and test computational models on a significantly larger dataset than those currently available, such as Ab-Bind [18] or SKEMPI.v2 [19], both of which contain less than 1k antibody-antigen specific data points. AbAgym will enable more accurate predictions of the impact of mutations on antibody-antigen affinity, thereby improving antibody design methodologies and enhancing variant escape prediction tools.

## Methods

### Data processing pipeline

All data in AbAgym were collected from scientific publications and preprints. Below, we summarize the main steps of its construction; for additional details, see Supplementary Materials.

#### Step 1: Collect and process the DMS data

First, we screened the literature and collected experimental DMS datasets of antibody-antigen complexes exclusively. The data was formatted as follows: for each single-site mutation, we recorded the wild-type and mutant residues, the protein chain in which the mutation occurs, the position of the mutated residue, and the experimental mutational score.

For homo-oligomeric antigens such as the HIV-1 envelope protein, mutations modify all chains, which is indicated in the “chains” column. All mutational scores have been checked to follow the usual sign convention of the Gibbs free energy, i.e. positive values correspond to a reduction in binding affinity. Values close to zero or negative values indicate affinity increase or absence of effect depending on the type of experimental assay (see Supplementary Material Section 1).

The majority of the entries in our dataset were downloaded in tabular form (hereafter referred to as “DMS table”) from the original paper. In some experiments, the DMS results were presented as a graphical heatmap, with each cell corresponding to a given mutation and colored according to its immune escape score. In these cases, we processed the heatmap image and mapped the RGB color of each cell to numeric values.

#### Step 2: Collect and process the PDB structure

The PDB codes for the 3D structures of antibody-antigen complexes used in the DMS studies were indicated in the original papers. They were used to interrogate the PDB. However, in the majority of these studies, the reported PDB structures exhibited minor differences from the DMS sequences, manifested by variations such as substitutions or missing residues or atoms. To address these discrepancies, we used the Swiss-Model algorithm [20] to remodel the structures to precisely match the mutagenesis sequence data. Finally, we performed energy minimization to relax the obtained structures and resolve possible atomic clashes, using the GROMACS suite with the Amber force field [21]. The resulting structures contain a single copy of the antibody, unlike the original PDBs which typically contain multiple copies.

The heavy and light chains of the antibodies were systematically renamed H and L, respectively, in the PDB files. Letters were assigned to the antigen chains, starting from A and in alphabetical order, and the residue numbering in the structures was adjusted to match exactly the residue numbers in the DMS tables.

#### Step 3: Identify the binding interface

The antibody-antigen binding interfaces were determined using a distance-based criterion. A residue is considered to be part of the antibody-antigen interface if at least one heavy atom in a residue of the antibody is at a distance ≤ 6 Å from at least one heavy atom in a residue of the antigen.

We have made all the collected DMS data available in a single table, where each row corresponds to a single-site mutation in a DMS study. The table includes the name of the associated modelled PDB file, the original PDB code used as a template, the mutated residue number, the wildtype and mutant residues, their distance from the binding interface, and the experimental DMS score. We have also made a table that provides metadata about each DMS dataset, including the names of the antibody and antigen chains, the range of the DMS score, whether the DMS was performed on the antigen or the antibody, the type of DMS experiment, and the DOI of the reference publication. AbAgym is freely available for academic use in our GitHub repository https://github.com/3BioCompBio/Abagym, including the data and metadata tables, and all the modelled PDB files.

## Results

### Data overview

AbAgym contains approximately 335,000 data points, including 37,361 interface mutations, each representing the impact of a single-site amino acid substitution on a given antibody-antigen complex. It includes different antigen types, with 30 datasets of the spike glycoprotein of the SARS-CoV-2 virus, especially its receptor-binding and N-terminal domains [22], 1 dataset of angiopoietin 2 [23], 15 of the HIV-1 envelope protein [24], 1 of the epidermal growth factor receptor (EGFR) [25], 4 of the influenza hemagglutinin [26], 1 of the lysosomal associated membrane protein 1 (LAMP-1) [27], 5 of Lassa virus’ glycoprotein complex (GPC) [28], 1 of the nerve growth factor (NGF) [29], 4 of the Nipah viral receptor-binding protein (RBP) [30], 1 of lysozyme [31], 1 of the vascular endothelial growth factor (VEGF) [32], and 3 of Zika’s viral envelope protein [33].

### DMS score type and distribution

The type of DMS scores we have collected in AbAgym vary across different DMS datasets, making it challenging to unify them. Indeed, AbAgym includes data obtained with different experimental techniques and mutational scores (see Supplementary Section 1 for details). All methods can test thousands of mutations in a single experiment, following similar but slightly different approaches [34, 35, 36, 37, 38]: site-directed mutagenesis allows the construction of a library of variants of a target antigen, which can be expressed either on the surface of yeast cells [34] or through a virion system [38, 35]. The variant library then undergoes a selection step by being exposed to neutralizing antibodies. The resulting mixture is subsequently sorted based on antibody binding in the case of yeast display, or used to infect target cells in an infection assay when using a pseudoviral system. Deep sequencing of the sorted mixture is then performed to quantify the frequency of each variant relative to a control condition. Note that similar approaches can also be applied to antibody-directed DMS experiments, where mutations are introduced into the antibody sequence [31].

The resulting distributions of DMS scores differ. The “escape fraction”, “antibody escape” and “fraction surviving” scores (Supplementary Section 1) range between zero and one, where zero indicates complete neutralization and one indicates complete escape. These scores are defined as the ratio between the frequency of a given variant in the antibody-treated condition and its frequency in the control, and typically show a unimodal distribution with a peak around zero, with just a few escaping mutations with scores close to one (Supplementary Figure 1.a).

In contrast, ‘differential selection’ and ‘enrichment ratio’ scores (Supplementary Section 1) range from negative to positive values, where zero indicates no effect on antibody binding, positive values indicate escape mutations, and negative values suggest enhanced binding. These scores are defined as the log ratio between the frequency of a given variant in the antibody-treated condition and its frequency in the control. They typically follow a unimodal distribution with approximately normal behaviour (Supplementary Figure 1.b).

Although DMS experiments provide a much more comprehensive and high-throughput approach than low-throughput binding assays, it also comes with some drawbacks. Being high-throughput, DMS is more susceptible to various sources of noise that can affect data quality. These include experimental sampling noise due to sequencing errors and depth, assay biases, cell line-specific effects, and variability between experimental replicates. For example, correlation coefficients between results from different experimental replicates can sometimes be relatively low.

### Structural analysis of hotspot residues

For structural analysis, we collected and curated 67 crystal structures in PDB format, one for each DMS dataset, each corresponding to a specific antibody bound to an antigen. When necessary, we remodelled the structures as described in the Methods section (step 2). The choice of the correct biological unit and of the conformation of each antigen is far from obvious. We describe and justify several cases where we made specific decisions, and give details about structure modeling and refinement in Supplementary Section 2. It is important to emphasize that the careful structural modeling we performed is essential for understanding the DMS scores and using them for, e.g., model training, as it allows the mutations to be accurately positioned within the correct structural context.

Interface residues with a strong impact on antibody-antigen binding affinity upon mutation, known as hotspot residues, are key to understanding phenomena such as immune evasion, which was critical, e.g., during the COVID-19 pandemic [22]. To identify and analyze hotspot residues in AbAgym, we calculated the average DMS score of all amino acid substitutions at each given position and selected the top five positions in each DMS dataset, which correspond to highly destabilizing positions upon mutation.

As expected, most hotspot positions are found to be close to the antibody-antigen interface, with 83% of them located within 6 Å of the interface (Figure 1.a). Mutations distant from the binding interface can induce long-range effects in the epitope or paratope regions, thereby indirectly affecting antibody–antigen binding. This has for example been shown in the case of the 12.1F antibody targeting the Lassa virus GPC, where the escape positions M134 and S135 are located at 13 Å and 18 Å away from the binding interface [15], respectively. In oligomeric antigen structures, mutations located at the oligomer interface can also alter the conformation of the assembly and lead to immune evasion. An example of this involves residues at positions D257 and R258 at the dimer interface of the receptor-binding domain of the Nipah virus, which are both located 20-30 Å away from the interaction site with the antibody, and whose mutation impairs recognition by the HENV-26 antibody [39]. In the case of the SARS-CoV-2 spike protein, the receptor-binding domain can adopt either an up or a down conformation, and therefore mutations that favor one of the two conformations could contribute to immune evasion by hiding specific epitopes, as previously shown [40, 41].

**Figure 1:**
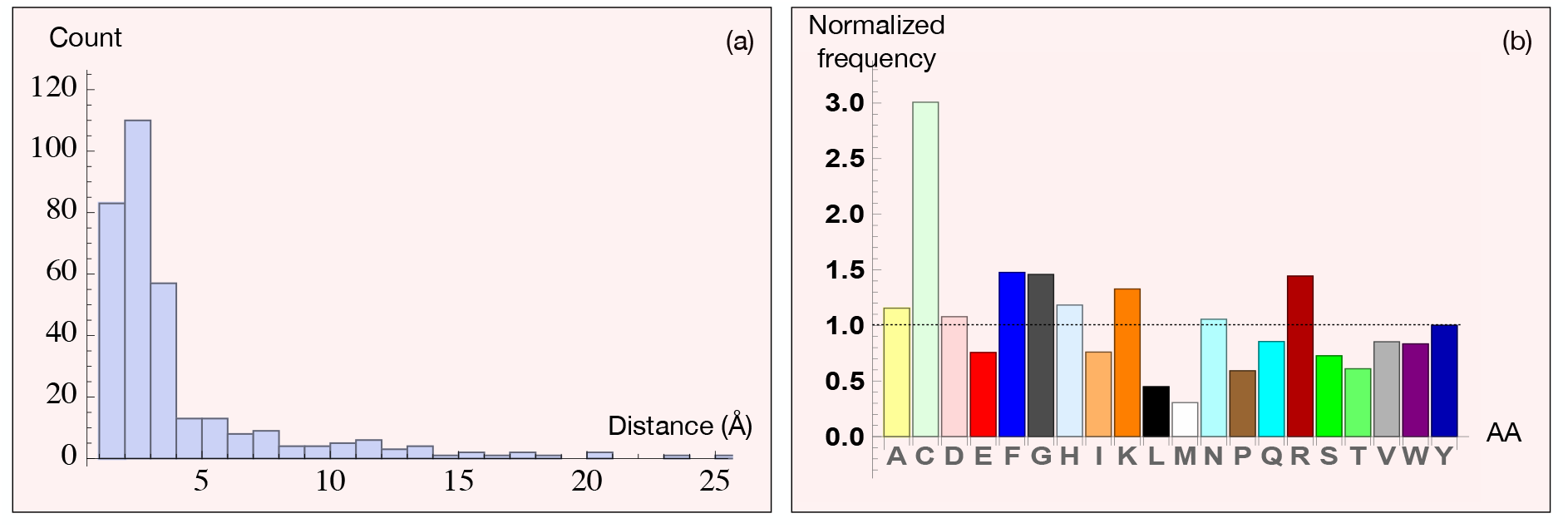
(a) Histogram (bin size = 1 Å) of the distance to the binding interface of all hotspot residues, which, as expected, shows that most hotspot residues are at the binding interface. (b) Amino acid frequency in the hotspots normalized by their frequency in the antibody-antigen interface.

These examples highlight the importance of accurately modelling the biological assembly and conformational state of antigen structures, which we carefully performed in our analysis. Indeed, both assembly and conformation can critically affect the interpretation of antibody binding.

We also analyzed the amino acid composition of interface hotspot residues to identify potential preferences. In Figure 1.b, this analysis reveals substantial variability in residue types, but without a strong overall trend. Notable exceptions are the enrichment of cysteine in hotspot positions, but also of phenyalanine, glycine and positively charged residues. Aromatic and positively charged residues are known to be important at the antibody-antigen interface, where they form *π*-*π* and cation-*π* interactions [3], and glycine is constrained likely because of structural flexibility requirements. Cysteine is rare at antibody-antigen interfaces but, when it appears, it often makes a disulfide bridge within the epitope and therefore clearly forms a hotspot. On the other hand, we observe a depletion of the hydrophobic residues leucine and methionine among the hotspots, suggesting that they are not critical for interface stability [3].

### Benchmarking of predictors

We evaluated a series of computational predictors on the 37,361 interface mutations in AbAgym. We tested six methods, grouped into three different classes: (1) Energy-based approaches (En), including FoldX [42], BeAtMuSiC [43] and korpm [44]; (2) evolutionary-based models (Evo), such as the log-odds ratio (LOR) [45] and the conservation index (CI), both of which measure conservation from the input multiple sequence alignment [45]; and (3) machine learning-based models (ML), including SAAMBE-3D [46], mCSM-AB [47], and SaProt [48], the latter using protein language models (pLMs). We used the residue solvent accessibility (RSA) computed as in [3] in the antibody-antigen complex as a baseline.

**Table 1:** Performance of different computational methods on the AbAgym dataset. We evaluated them using the Spearman correlation coefficient (*ρ*) and the ROC-AUC, where the experimental values were binarized as described in the text.

| Method | Type | Average Spearman $\rho$ | Average ROC-AUC |
| --- | --- | --- | --- |
| FoldX [42] | En | 0.25 | 0.72 |
| BeAtMuSiC [43] | En | 0.20 | 0.62 |
| korpm [44] | En | 0.04 | 0.49 |
| CI [45] | Evo | -0.02 | 0.44 |
| LOR [45] | Evo | 0.03 | 0.48 |
| SAAMBE-3D [46] | ML | 0.11 | 0.59 |
| mCSM-AB [47] | ML | 0.01 | 0.49 |
| SaProt [48] | pLM | 0.01 | 0.54 |
| RSA | Baseline | 0.28 | 0.68 |

As performance metrics, we used the Spearman correlation coefficient between experimental and predicted values, and the area under the receiver operating characteristic curve (ROC-AUC). For the latter, experimental values were binarized by labeling the top 5% most destabilizing mutations in each DMS as ‘destabilizing’, and the remaining as ‘neutral’, following a procedure similar to that described in [49].

Overall, we found that the Spearman correlation coefficients and ROC-AUC values between experimental and predicted results are modest (with maxima of 0.28 and 0.72, respectively), and that even the best-performing methods achieve performances comparable to the RSA baseline. This underscores the fact that despite impressive performances in cross validation often reported in the current literature [50], accurately predicting the impact of mutations on antibody-antigen binding affinity remains an open challenge. A detailed comparison for each of the targets can be found in Supplementary Tables 1 and 2.

Energy-based methods are the most robust, with FoldX slightly outperforming the others. In contrast, both evolutionary models LOR and CI perform poorly, which is expected, as in antibody–antigen complexes, evolutionary dynamics play a intricate role. Finally, machine learning models also fail to produce statistically meaningful predictions, which suggests they have been overtrained and do not generalize outside their training set, unlike general energy-based methods. This result is in agreement with an independent, recently published, antibody-antigen mutational dataset whose authors also reported that recent ML models perform much worse than the FoldX force field [50]. Interestingly, we observe a correlation between the interface size and the performance of the three energy-based methods: mutations in larger interfaces are generally predicted less accurately than mutations in smaller interfaces.

## Conclusion

AbAgym is the first large-scale, diverse and carefully curated mutational dataset specific to antibody-antigen interactions. By making AbAgym freely available for academic use, we aim to support the training, evaluation, and improvement of current computational methods to predict the impact of mutations on antibody-antigen binding affinity, which is one of the cornerstones of in silico antibody optimization.

## Acknowledgments

We thank the FNRS-Belgian Fund for Scientific Research for financial support through a PDR project; D.L. benefits from a scholarship from the China Scholarship Council.

