## Supplementary Material for "AbAgym: a well-curated dataset for the mutational analysis of antibody-antigen complexes"

Gabriel Cia[1], Dong Li[1], Simon Poblete[3], Marianne Rooman[1,2] and Fabrizio Pucci[1,2]
[1]*Computational Biology and Bioinformatics, Université Libre de Bruxelles, 1050 Brussels, Belgium*
[2]*Interuniversity Institute of Bioinformatics in Brussels, 1050, Belgium*
[3]*Facultad de Ingeniería, Arquitectura y Diseño, Universidad San Sebastián, Santiago, 7500912, Chile*

**Index**

1. Experimental methods
2. Structural modeling and refinement
3. Predictors' performance tables

### 1 Experimental Methods

We provide a description of the experimental setups used in the different DMS datasets included in AbAgym:

- **Escape Ratio**. This technique combines yeast cells that express on their surface fluorescently tagged single-site variants of the SARS-CoV-2 receptor-binding domain (RBD) in combination with a fluorescently tagged antibody of interest. Fluorescence-activated cell sorting (FACS) is then used to first filter variants that impair proper protein folding or binding to the ACE2 receptor, ensuring that only structurally and functionally viable mutants are analysed, followed by a second round of FACS to quantify the fraction of RBD-expressing cells that escape antibody binding. This escape fraction reflects the extent to which a mutation disrupts antibody recognition: values close to 1 indicate a strong escape (i.e., the mutation abolishes antibody binding), whereas values close to 0 suggest no measurable effect on antibody interaction [1].
- **Enrichment ratio.** Site-saturation mutagenesis is used to generate a comprehensive library of mutant proteins, which are then displayed on the surface of yeast cells. The cells are exposed to a neutralizing antibody, and the impact of each mutation is quantified using an enrichment ratio score. This score is calculated as the $\log_2$ ratio between the frequency of yeast cells expressing a given mutation in the antibody-sorted population and its frequency in an unsorted (reference) population. A negative score indicates that the mutation enhances antibody binding (i.e., increases affinity), while a positive score indicates that the mutation reduces binding, suggesting immune escape. Note that a similar approach can also be applied to antibody-directed DMS experiments, where mutations are introduced into the antibody sequence, typically in the complementary-determining regions [2].
- **Antibody Escape.** First, a pseudovirus variant library is generated, with each pseudovirus expressing a single amino acid variant of the viral antigen. This library is then exposed to neutralizing antibodies at varying concentrations. The antibody–virus mixtures are subsequently used in infection assays to assess the ability of each variant to infect target cells. The antibody escape score quantifies the relative abundance of each variant in the presence of antibodies compared to the antibody-free control, normalized using a neutralization standard [3]. Escape scores range from 0 (indicating complete neutralization by the antibody) to 1 (indicating complete escape from neutralization) [3].
- **Mutational differential selection.** First, a library of viral clones is generated, with each clone encoding a unique single amino acid variant of the target protein. The library is initially screened in an infection assay to ensure that only functional variants are retained. The selected library is then exposed to a specific neutralizing antibody as well as to a positive control antibody. The resulting antibody–virus mixtures are used to infect target cells. To assess the effect of each mutation, viral cDNA is extracted from the infected cells, and the frequency of each variant is measured. The differential selection score is calculated as the log ratio of a mutation's frequency (normalized to wild-type) in the antibody-treated condition versus the control. Mutations with positive differential selection scores are enriched in the presence of the neutralizing antibody, indicating escape, while those with negative scores suggest increased sensitivity to neutralization [4, 5].
- **Mutational fraction survival.** The experimental procedure follows the same approach as mutational differential selection. However, in this case, the scores are calculated as the fraction of each mutation observed in the antibody-treated sample relative to the control, with corrections applied to account for sequencing errors and differences in read depth [6]. These values range from 0, indicating complete neutralization by the antibody, to 1, indicating complete escape from neutralization.

The distribution of the DMS scores varies depending on the type of assay. In particular, assays such as the "Escape Ratio", "Mutational Fraction Survival" and "Antibody Escape", typically produce distributions that are bounded between 0 and 1, with a prominent peak near zero and a smaller secondary peak corresponding to immune escape variants (see an example in Figure 1a). In contrast, assays based on "Enrichment Ratio" or "Mutational Differential Selection" exhibit unimodal distributions centered around zero (see an example in Figure 1b).

### 2 Structural Modeling and Refinement

The choice of the correct biological unit and of the conformation of each antigen is far from trivial. Here, we describe and justify several cases in AbAgym where specific decisions were made. In the first eight antigens described,

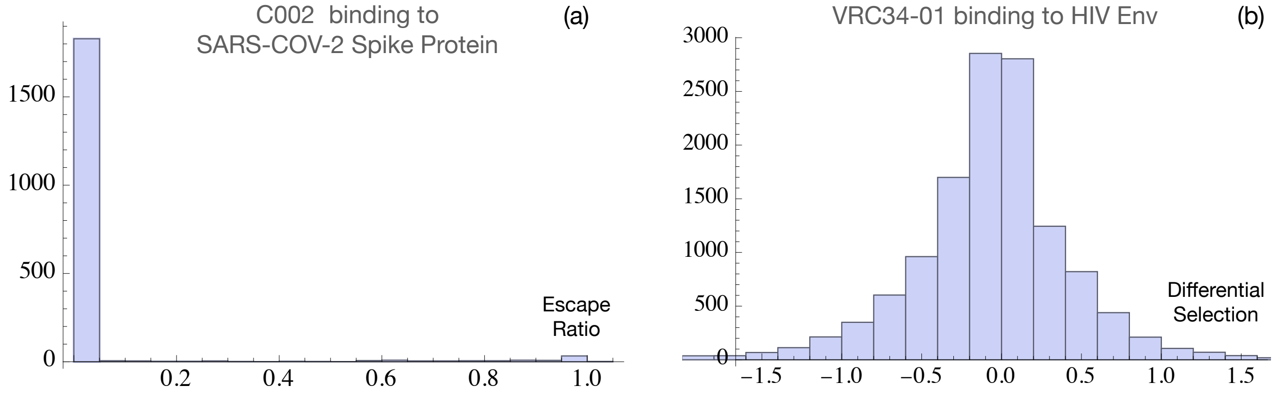

Figure 1: Examples of score distributions for two DMS datasets: (a) the escape ratio scores for the binding of the C002 antibody to the SARS-CoV-2 spike protein, and (b) the mutation differential selection scores for the binding of the VRC34-01 antibody to the HIV Env protein.

mutagenesis was performed on the antigen residues in the DMS experiment, and in the last four, it was performed on the antibody residues.

**Nipah RBP**: The receptor-binding protein (RBP) of the Nipah virus, also known as the NiV G protein or NiV G ectodomain, is a homotetramer with each subunit composed of three domains (see Figure 2a): a stalk domain that mediates tetramerization and is characterized by a long $\alpha$-helix; a neck domain composed of short $\beta$-strands; followed by a globular head domain. The linker between the neck and the head domain is highly flexible, making two head domains fold back toward the viral membrane, thus proximal to the viral membrane and distant from the receptor-binding region, while the other two head domains remain in an upright position, thus more accessible to the host receptor [7].

In our dataset, four RBP-antibody complexes are included. Two of these antibodies, HENV26 and HENV32, are neutralizing antibodies originally developed against the Hendra virus. Due to the high similarity between the epitopes of the Hendra virus and the Nipah virus, these antibodies were used in the DMS experiment to neutralize the Nipah virus.

Two of the four antibodies (HENV26 and nAH1.3) are neutralizing and bind to epitopes that are accessible in the context of the full tetrameric RBP. We thus included the two host-receptor-proximal head domains in the PDB structure to study residues at the interface between the two head domains, which can potentially be the hotspot residues affecting antibody-antigen binding despite being far from the paratope. The other two antibodies bind to epitopes on the head domain; however, these two antibodies were developed using only the head domain as the antigen, and our structural analysis showed that they would clash with other subunits when docked onto the full tetramer, suggesting that their epitopes are inaccessible in this conformation of the tetrameric complex. For these two complexes, we only included one head domain in each PDB file as the antigen.

**HIV envelope (Env) protein**: HIV mainly infects immune cells that express CD4 and co-receptors such as CCR5 and CXCR4. The HIV envelope glycoprotein (Env) is the main protein responsible for cell entry. Env is a homotrimer, with each protomer consisting of two subunits: gp120 and gp41, both derived from a gp160 precursor. Binding of gp120 to CD4 induces a conformational change that exposes the co-receptor binding site. Subsequent engagement with the co-receptor (e.g., CCR5 or CXCR4) triggers further conformational changes in gp41, promoting membrane fusion between the viral and host cell membranes. Two distinct binding interfaces exist on gp120 – one for CD4 and another for the co-receptor. Thus, membrane fusion involves a two-step triggering mechanism: CD4 binding primes gp120 by exposing the co-receptor binding site, and co-receptor binding activates gp41-mediated fusion [8]; see Figure 2c for an example of the structure of HIV Env bound to the antibody 3BN-1074.

In our dataset, we modeled the trimeric structure of the Env protein in Env-antibody complexes when the correct structure was unavailable. In cases where the sequence used for modeling was derived from wild-type residues in DMS experiments, but the chain assignment (gp120 or gp41) for each residue was not labeled, we manually assigned the chains to derive separate sequences for gp120 and gp41. The three gp120 chains were named A, C, and E; and the gp41 chains were named B, D, and F.

**Influenza HA**: Influenza viruses are classified into types A, B, C, and D, depending on the glycoproteins expressed on their surface. Influenza C causes mild symptoms in humans, and influenza D has not been reported to infect humans. Influenza A and B viruses cause more severe symptoms and spread more rapidly, with Influenza A capable of triggering pandemics. As a result, they have attracted greater scientific attention and have been studied more extensively than C and D. Surface glycoproteins on influenza A and B viruses are HA and neuraminidase (NA), both of which can be targeted by antibodies. Cell entry of influenza A and B viruses is mediated by the HA binding to host receptors. Influenza A viruses are further classified into subtypes based on HA and NA, with 18 types of HAs (H1-H18) and 11 subtypes of NAs (N1-N11) [9]. In our dataset, we included four HA-antibody complexes, all from influenza A viruses. These are: antibodies C179, S139 and FI6v3 bound to H1N1 influenza HAs, and FI6v3 bound to a H3N2 influenza HA.

HA is a homotrimer in which each subunit is composed of two non-covalently-linked domains: a membrane-distal globular head domain responsible for receptor binding, on top of a membrane-proximal stem domain (see Figure 2d). Despite the fact that the stem domain does not directly engage with the host receptor, the structural rearrangement of this domain also plays an indispensable role in membrane fusion of influenza viruses, making the stem domain a useful target for broadly neutralizing antibodies as well. In our dataset, C179 and FI6v3 are stem-targeting antibodies and S139 targets the head domain.

For the four complexes, the HA sequences in the original PDBs are not the same as those derived from the DMS experiments. We therefore remodeled the four structures using the DMS sequences and the original PDBs as templates. The three head domains were named A, C and E; the three stem domains were named B, D and F.

**Lassa GPC**: The glycoprotein complex (GPC) mediates the entry of the Lassa virus into the host cell. It is the main surface protein displayed by Lassa viruses and the sole target of neutralizing antibodies. The Lassa GPC is a homotrimer and each subunit consists of three domains: a stable signal peptide (SSP), followed by the glycoprotein 1 (GP1) domain, and the glycoprotein 2 (GP2) domain which non-covalently associates with SSP and GP1. SSP is composed of two hydrophobic $\alpha$-helices (denoted h1 and h2), among which h1 is transmembrane. GP1 is the receptor binding domain and GP2 is the membrane proximal domain responsible for the fusion between host and viral membranes [10, 11]. Both GP1 and GP2 contain epitopes targeted by neutralizing antibodies. In our dataset, we included five Lassa GPC-antibody complexes: three in which the antibody is bound to the GP2 subunit and two in which it is bound to GP1. Figure 3a-c shows an example of the structure of Lassa GPC bound to antibody 377H.

In the DMS experiment, the sequences of the antigens in the five complexes are the same, but the sequences in the reference PDBs differ from the DMS sequences by several residues. We therefore remodeled the complexes. The three GP1 domains were named A,C and E; the three GP2 domains B, D and F.

**Zika E protein dimer**: The particle of a mature Zika virus that induces host-cell membrane fusion is the E protein. Two E proteins form a homodimer and each E protein has three domains: the N-terminal DI domain with a transmembrane $\alpha$-helix-rich region, the DII domain responsible for dimerization and the C-terminal DIII domain with an immunoglobulin fold. The Zika virus surface contains 180 E protein monomers (90 dimers). These dimers are further organized into 30 groups, arranged side by side in each group. Each group is conventionally referred to as a "raft", and together, the 30 rafts uniformly distribute over the viral surface. Additionally, two small transmembrane M proteins form a complex with the E protein dimer, each accompanying the transmembrane region of the DI domain of each E protein in the dimer. Figure 4a–c illustrate the structure of Zika virus' E protein dimers and a complex in which the E dimer is bound to the EDE1-C10 antibody.

AbAgym contains three Zika antibody-antigen complexes. These are the complexes with antibody ZV-67, MZ4 and EDE1-C10, respectively. All these antibodies bind to the same antigen. However, the reference PDBs of these three complexes contain problematic structures; here is how we treated them:

- The antigen in the PDB of EDE1-C10 misses 10 residues covered in the DMS experiment. We thus remodeled the antigen with the DMS sequence and removed the two chains of M proteins as they are not in contact with the antibody.
- The antigen in the PDB of the ZV-67 complex is a small fragment of the E protein. To complete it, we directly

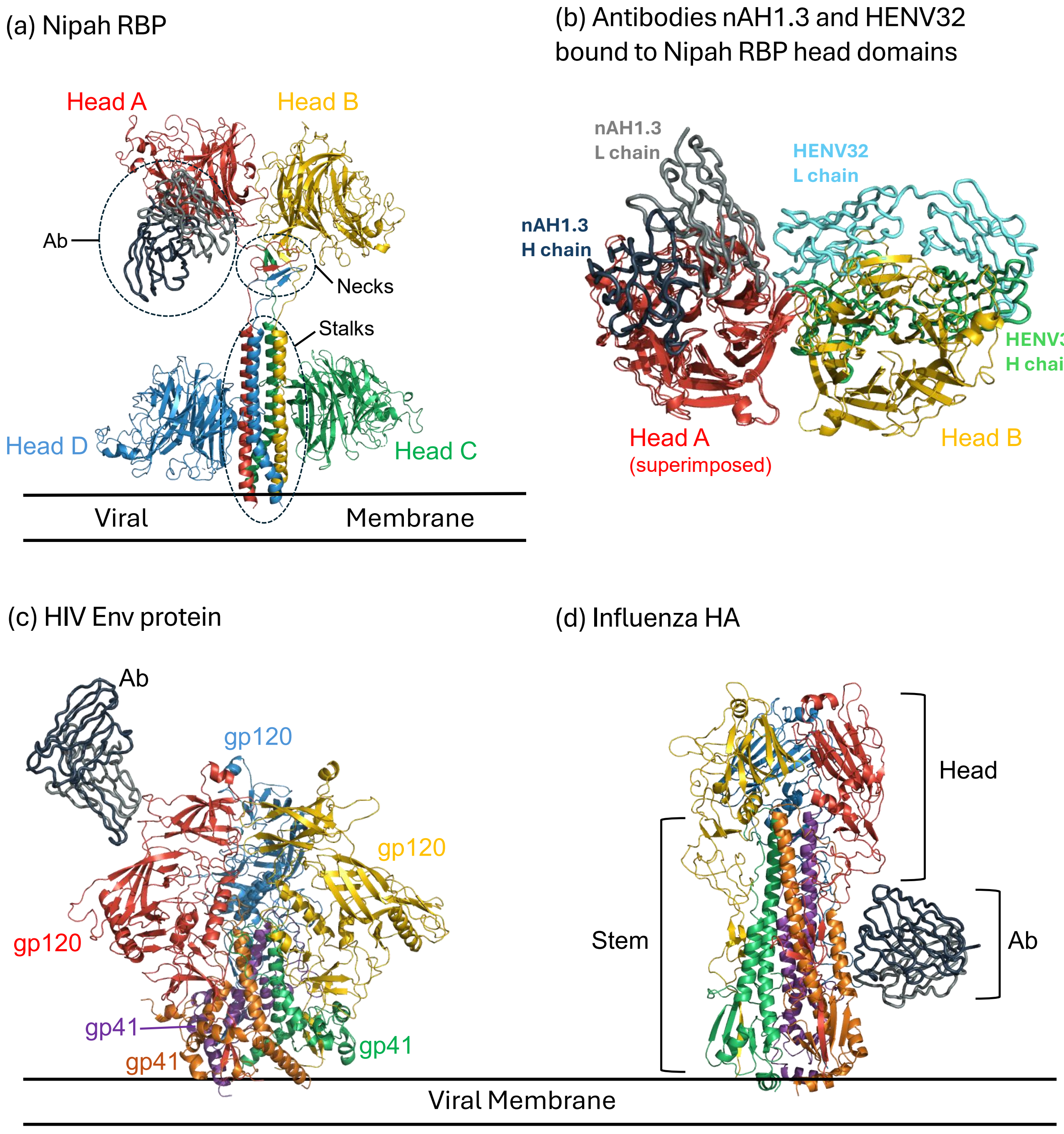

Figure 2: Structures of three viral antigens bound to antibodies: Nipah viral receptor-binding protein (RBP) (a-b), HIV Env protein (c) and influenza viral hemagluttinin (HA) (d). The heavy chains of the antibodies are shown in dark blue (and green in panel (b) for antibody HENV32) and light chains in grey (and cyan in panel (b) for antibody HENV32). The membrane is represented schematically as two parallel lines labeled "viral membrane". (a) Structure of the Nipah RBP anchored to the viral membrane, bound to antibody nAH1.3. The four identical subunits from A to D are colored in red, yellow, green and blue, respectively. (b) Two RBP-antibody complexes, involving the antibodies nAH1.3 and HENV32, respectively, superimposed on the head domain A (in red). The antibody nAH1.3 can bind the whole tetrameric RBP structure without steric clashes (obviously it does not clash with the head domain B (in yellow)). In contrast, HENV32 can only access the single free head domain A and it would sterically clash with domain B in the context of the full RBP tetramer. (c) Structure of the HIV Env protein anchored to the membrane, bound to antibody 3BN-1074. The three gp120 subunits are colored in red, yellow and blue, respectively; and the three gp41 subunits, in orange, green and purple. (d) Structure of influenza HA anchored to the membrane, bound to antibody FI6V3-H3. The three head domains are colored in red, yellow and blue, respectively; and the three stem domains, in orange, green and purple.

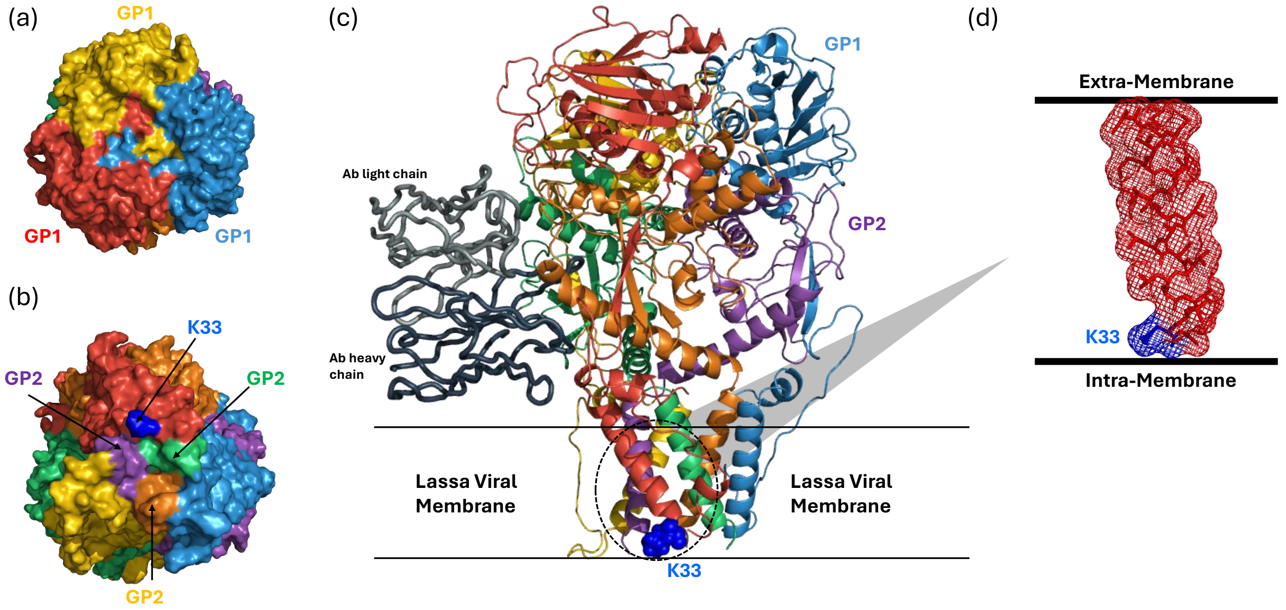

Figure 3: Structural description of the Lassa viral glycoprotein complex (GPC). Residue Lys33 is shown in light blue and labeled "K33". (a) Top view of GPC. The three GP1 domains are colored in red, yellow and blue, respectively. (b) Bottom view of GPC. The three GP2 domains are colored in orange, green and purple, respectively. (c) Structure of GPC bound to antibody 377H. The heavy and light chains of the antibody are colored in dark blue and grey, respectively. The colors of the six GPC domains are the same as in panels (a) and (b). (d) The structure of the SSP transmembrane domain and residue Lys33, which is located near the inner viral membrane.

took the abovementioned remodeled structure and superimposed it onto the fragment, followed by molecular dynamics relaxation.

- In the PDB of the MZ4 complex, the antigen misses coordinates of several residues. We likewise superimposed the remodeled structure onto the original one.

**SARS-CoV-2 spike protein**: Infection by SARS-CoV-2 is mediated by the fusion of the viral membrane with the host cell's membrane, through binding of the spike (S) glycoprotein with the angiotensin-converting enzyme 2 (ACE2) at the host cell's surface. The S protein forms a homotrimer with each monomeric subunit containing, in order from the N- to the C-terminus, an N-terminal domain (NTD), an RBD which binds ACE2 and is the principal target of many neutralizing antibodies, and subdomains 1 and 2 (SD1 and SD2) [12].

The RBD has two conformations: the up and down conformations, and only the up conformation exposes the receptor-binding site to ACE2, meaning that the activation of the trimeric spike protein requires the RBD of at least one S monomer to adopt the up conformation (see Figure 5a-b for the structure of the spike protein with its RBD in different conformations (a) and bound to antibody C121 (b-c)).

AbAgym includes 30 RBD-antibody complexes. In eight of them, the DMS experiments were performed on the full-length spike monomer, while in the remaining twenty-two, only the RBD region was assessed. For the former, we remodeled the antigen as the spike trimer; for the latter, we remodeled it as the RBD.

**$\beta$-NGF**: $\beta$-NGF naturally forms a homodimer, whereas the antigen in the reference PDB contains only a monomer. We therefore remodeled and dimerized the antigen to reflect its native state.

**LAMP-1**: Human LAMP-1 is a transmembrane protein. In the DMS experiment, only its extramembrane domain was studied. This was in part because in the experiment the protein was expressed on the surface of yeast cells rather than human cells, which was made feasible by replacing the native transmembrane domain of LAMP-1 with a yeast cell surface anchor. Consequently, only the extramembrane domain retained its native sequence and structure [13].

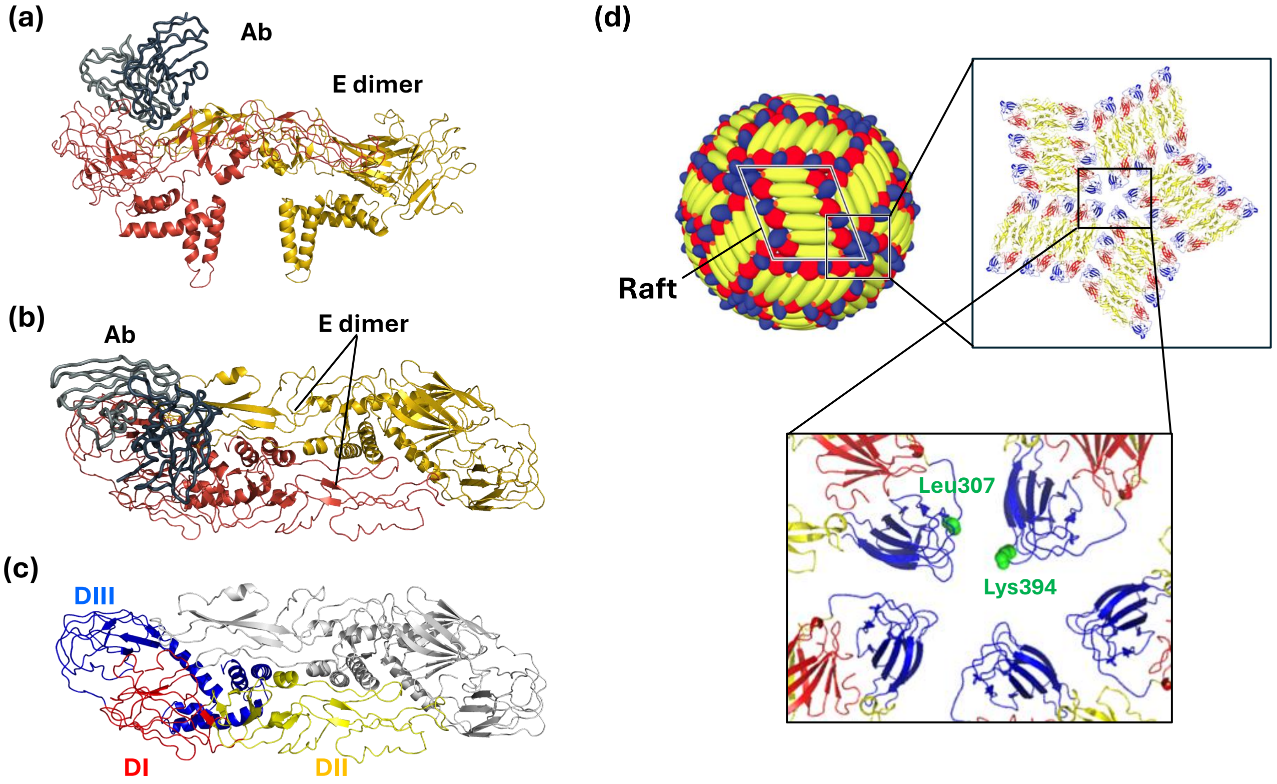

Figure 4: Structural description of the Zika virus E protein. (a) Side view of the E protein dimer bound to the antibody EDE1-C10. The two chains of the E dimer are shown in red and yellow, respectively. (b) Top view of panel (a). (c) Domain organization of an E monomer: domains DI, DII, and DIII are colored red, yellow, and blue, respectively. The second monomer in the dimer is shown in grey. (d) Schematic view of rafts on the surface of the Zika virus. A pentagon formed by the head-to-head arrangement of five rafts is shown. Two residues, Lys394 and Leu307, located on different rafts, are highlighted in green; their inter-raft interaction is believed to stabilize the overall raft conformation.

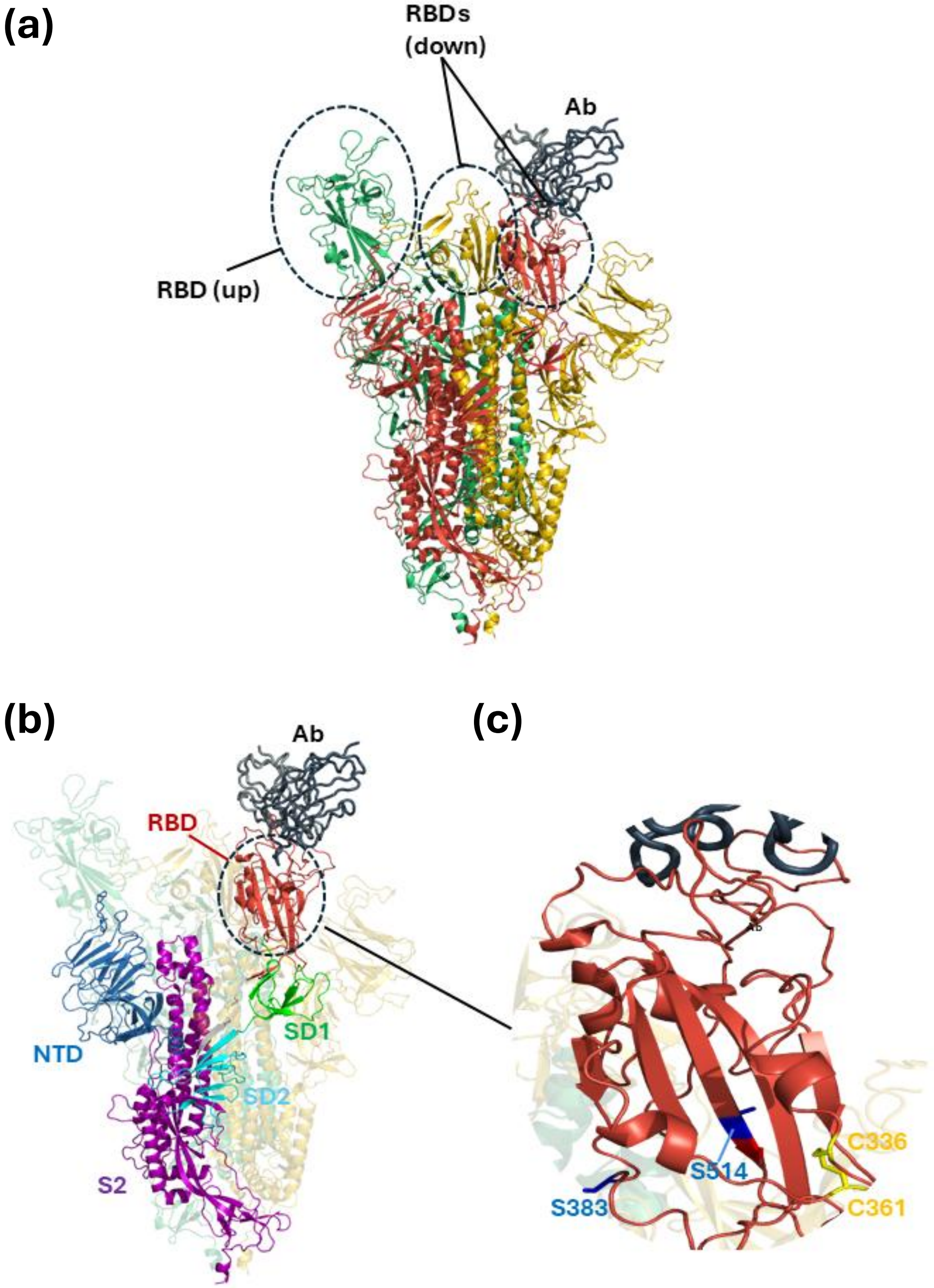

Figure 5: Structural description of the SARS-CoV-2 spike protein bound to antibody C121. The heavy and light chains of the antibody are colored in dark blue and grey, respectively. (a) Structure of the spike protein bound to antibody C121. The three chains of the spike protein are colored in red, yellow and green, respectively. (b) Domains in one S protein: the N-terminal domain (NTD, in blue), the receptor-binding domain (RBD, in red), subdomain 1 (SD1, in green) and subdomain 2 (SD2, in cyan), and the S2 subunit (in purple). (c) Highlighted are the two cysteine residues (Cys336 and Cys361, in yellow) that form a disulfide bond and two serines (Ser383 and Ser514, in blue) involved in the regulation of the RBD conformation.

For the following antigens, the mutagenesis experiments were performed on the antibody residues. We will not go into details of the structures of these antigens, but provide a brief description of these proteins and associated antibodies.

**Lysozyme**: Lysozyme is a small, monomeric, globular antimicrobial enzyme. It is widely distributed in bodily secretions and immune cells such as macrophages, where it helps destroy bacterial cell walls. In this DMS experiment, mutations were introduced into antibody D44.1, which targets hen egg white lysozyme, to identify hotspot residues contributing significantly to antibody-antigen complex stabilization [14].

**EGFR**: EGFR is a transmembrane receptor involved in regulating cell growth and proliferation. Since the antibody used in this study targets the extracellular portion of EGFR, only this region was retained and reconstructed as the extracellular domain. The antigen was expressed in solution, while the mutant antibodies were expressed and displayed on the surface of mammalian cells [15].

**Angiopoietin-2**: Angiopoietin-2 is one of the four types of angiopoietins, secreted glycoproteins involved in angiogenesis and vascular remodeling. It contains a coiled-coil domain responsible for oligomerization and a fibrinogen-like domain that binds the Tie2 receptor on endothelial cells.

The antibody used in this DMS experiment was a dual-action antibody with dual specificity for angiopoietin-2 and the vascular endothelial growth factor (VEGF, discussed below). Two rounds of separate DMS experiments were performed on the same antibody: the first round against VEGF, and the second round against angiopoietin-2. The goal was to identify key hotspot residues that contribute primarily to binding stability with each antigen, respectively. These residues would then be targeted for mutation in order to design a paratope capable of simultaneously binding both VEGF and angiopoietin-2, creating an antibody with dual-action antigen-binding region [16].

**VEGF**: VEGF is a dimeric signaling protein that stimulates angiogenesis by binding VEGF receptors on endothelial cells. In the reference antibody-antigen structure, the antigen is monomeric; we did not remodel it to build a dimer because the DMS data is specific to the antibody. The antibody was discussed above [16].

### 3 Predictors' Performance Tables

| DMS DATA | # mut | ACC | BeAt MuSiC | FoldX | CI | LOR | mCSM AB | SAAMBE 3D | SaProt | Korpm |
|---|---|---|---|---|---|---|---|---|---|---|
| 1-18_6udj | 690 | 0.307 | 0.137 | 0.210 | 0.070 | 0.049 | 0.070 | 0.104 | 0.065 | 0.013 |
| 106E6_6n1w | 551 | 0.080 | 0.193 | 0.170 | -0.390 | -0.234 | 0.150 | -0.209 | 0.070 | -0.002 |
| 121F_7uov | 482 | 0.099 | 0.182 | 0.120 | 0.128 | 0.189 | -0.070 | 0.022 | -0.068 | 0.019 |
| 17D4_6n1v | 475 | 0.527 | 0.270 | 0.140 | -0.415 | -0.387 | 0.340 | 0.003 | 0.130 | -0.007 |
| 256A_6p95 | 1257 | 0.129 | 0.024 | 0.130 | 0.015 | 0.188 | 0.000 | 0.044 | 0.034 | 0.042 |
| 372D_7uot | 967 | 0.090 | 0.068 | -0.010 | 0.042 | 0.049 | 0.000 | -0.001 | -0.100 | 0.017 |
| 377H_5vk2 | 1346 | 0.173 | 0.089 | 0.100 | 0.063 | 0.031 | -0.030 | 0.121 | -0.030 | 0.041 |
| 3BN-1074_5t3z | 475 | 0.222 | 0.184 | 0.260 | 0.240 | 0.149 | 0.020 | 0.097 | 0.073 | 0.009 |
| 89F_7uot | 731 | 0.186 | 0.167 | 0.220 | 0.177 | 0.152 | -0.150 | 0.181 | 0.007 | 0.005 |
| AZD1061_7l7e | 395 | 0.321 | 0.225 | 0.400 | 0.124 | 0.130 | -0.020 | 0.218 | -0.011 | -0.040 |
| AZD8895_7l7d | 345 | 0.396 | 0.322 | 0.340 | -0.083 | 0.067 | -0.210 | 0.298 | -0.036 | 0.037 |
| BD55-5840_7wrz | 88 | 0.125 | 0.142 | 0.240 | 0.199 | 0.155 | -0.130 | 0.107 | -0.142 | 0.058 |
| C002_7k8s | 578 | 0.431 | 0.291 | 0.360 | 0.001 | 0.041 | 0.080 | 0.307 | 0.145 | 0.055 |
| C105_6xcm | 430 | 0.459 | 0.261 | 0.040 | 0.183 | 0.109 | -0.040 | 0.009 | 0.207 | 0.018 |
| C110_7k8v | 345 | 0.154 | 0.309 | 0.180 | -0.156 | -0.004 | 0.020 | 0.203 | 0.024 | 0.091 |
| C119_7k8w | 243 | 0.168 | 0.191 | 0.090 | 0.118 | 0.060 | 0.050 | 0.166 | 0.049 | 0.150 |
| C121_7k8x | 642 | 0.449 | 0.511 | 0.290 | -0.171 | -0.028 | -0.210 | 0.398 | -0.178 | -0.114 |
| C135_7k8z | 220 | 0.304 | 0.059 | 0.410 | -0.155 | -0.020 | -0.220 | 0.369 | 0.243 | -0.018 |
| C144_7k90 | 554 | 0.385 | 0.316 | 0.300 | -0.246 | -0.114 | 0.050 | 0.239 | -0.009 | -0.004 |
| C179_4hlz | 855 | 0.222 | 0.096 | 0.060 | 0.144 | 0.058 | -0.010 | 0.095 | 0.069 | 0.089 |
| COV2-2130_8d8q | 650 | 0.439 | 0.298 | 0.610 | 0.115 | 0.179 | -0.040 | 0.403 | 0.067 | 0.151 |
| COV2-2196_8d8q | 495 | 0.215 | 0.312 | 0.300 | -0.062 | 0.094 | -0.250 | 0.215 | 0.016 | 0.137 |
| COVA2-04_7jmo | 292 | 0.214 | 0.214 | 0.280 | 0.160 | 0.133 | 0.140 | 0.129 | 0.192 | 0.114 |
| CR3022_6zlr | 456 | 0.327 | 0.362 | 0.300 | 0.027 | 0.035 | -0.080 | 0.225 | -0.123 | -0.025 |
| Cetuximab_1yy9 | 659 | 0.346 | 0.083 | 0.210 | -0.366 | -0.122 | -0.060 | 0.300 | 0.092 | 0.095 |
| D441_1mlc | 627 | 0.250 | 0.157 | 0.170 | 0.007 | 0.108 | -0.060 | 0.171 | -0.126 | 0.064 |
| DF1W314_6mph | 722 | 0.241 | 0.096 | 0.230 | -0.289 | -0.186 | -0.090 | -0.167 | 0.072 | 0.011 |
| EDE1-C10_5h37 | 1501 | 0.166 | 0.045 | 0.040 | -0.012 | 0.020 | -0.050 | -0.063 | -0.153 | -0.020 |
| FAB-B_8ath | 399 | 0.363 | 0.281 | 0.340 | 0.103 | 0.139 | 0.060 | 0.119 | -0.204 | 0.084 |
| FI6V3-H3_3ztj | 513 | -0.016 | 0.142 | 0.050 | 0.011 | 0.077 | 0.070 | 0.253 | -0.056 | -0.024 |
| FI6V_3ztn | 703 | 0.337 | 0.056 | 0.130 | 0.146 | -0.097 | 0.020 | -0.096 | 0.057 | 0.063 |
| FP16-02_6cdi | 608 | 0.405 | 0.220 | 0.270 | -0.305 | -0.119 | 0.150 | -0.166 | 0.111 | 0.003 |
| FP20-01_6cde | 665 | 0.295 | 0.283 | 0.170 | -0.364 | -0.149 | 0.130 | -0.104 | 0.190 | 0.001 |
| G6_27_30A_4zff | 475 | 0.461 | 0.088 | 0.260 | 0.145 | 0.016 | -0.060 | -0.168 | -0.185 | 0.069 |
| G6_27_30A_4zfg | 471 | 0.414 | 0.211 | 0.370 | 0.148 | 0.138 | 0.160 | 0.047 | -0.321 | 0.106 |

| DMS DATA | # mut | ACC | BeAt MuSiC | FoldX | CI | LOR | mCSM AB | SAAMBE 3D | SaProt | Korpm |
|---|---|---|---|---|---|---|---|---|---|---|
| HENV26_6vy5 | 1144 | -0.205 | -0.141 | 0.020 | -0.038 | -0.041 | 0.000 | -0.028 | -0.011 | 0.001 |
| HENV32_6vy4 | 814 | -0.073 | -0.071 | 0.000 | -0.296 | -0.156 | 0.030 | 0.038 | -0.073 | -0.025 |
| LY-CoV016_7c01 | 510 | 0.457 | 0.321 | 0.340 | 0.212 | 0.169 | 0.050 | 0.079 | 0.111 | -0.033 |
| LY-CoV1404_7mmo | 170 | 0.076 | 0.266 | 0.170 | -0.242 | -0.152 | 0.170 | 0.175 | -0.022 | 0.228 |
| LY-CoV488_7kmh | 207 | 0.347 | 0.231 | 0.450 | 0.068 | 0.236 | -0.030 | 0.080 | -0.050 | -0.051 |
| LY-CoV555_7kmg | 431 | 0.471 | 0.352 | 0.460 | -0.397 | -0.216 | 0.110 | 0.358 | 0.053 | 0.096 |
| MZ4_6niu | 741 | 0.256 | 0.145 | 0.230 | 0.098 | 0.136 | 0.140 | 0.093 | -0.052 | -0.006 |
| OPV12_6ot1 | 532 | 0.386 | 0.165 | 0.210 | -0.348 | -0.011 | 0.320 | -0.276 | 0.157 | 0.023 |
| OPV20_6osy | 570 | 0.267 | 0.281 | 0.040 | -0.530 | -0.263 | -0.010 | -0.176 | 0.192 | -0.005 |
| PG9_3u4e | 437 | 0.273 | 0.020 | 0.110 | -0.031 | 0.086 | -0.070 | 0.250 | 0.083 | -0.017 |
| PGT121_7uoj | 399 | 0.221 | 0.172 | 0.370 | 0.304 | 0.277 | 0.040 | 0.011 | 0.099 | 0.007 |
| PGT145_5v8l | 646 | -0.148 | 0.029 | 0.340 | -0.072 | -0.001 | -0.270 | 0.027 | 0.079 | 0.010 |
| PGT151_5fuu | 817 | 0.231 | 0.020 | 0.170 | -0.087 | -0.008 | -0.100 | -0.003 | -0.030 | 0.021 |
| REGN10933_6xdg | 385 | 0.432 | 0.368 | 0.490 | 0.005 | 0.111 | -0.030 | 0.273 | 0.209 | -0.002 |
| REGN10987_6xdg | 265 | 0.672 | 0.443 | 0.310 | -0.129 | 0.133 | -0.010 | 0.164 | -0.085 | 0.051 |
| S139_4gms | 741 | 0.263 | 0.022 | 0.110 | 0.062 | -0.078 | 0.030 | 0.061 | -0.083 | 0.001 |
| S2D106_7r7n | 470 | 0.272 | 0.381 | 0.390 | -0.099 | -0.096 | -0.010 | 0.319 | 0.263 | 0.122 |
| S2E12_7r6x | 341 | 0.548 | 0.495 | 0.640 | -0.164 | 0.055 | -0.300 | 0.399 | 0.043 | 0.130 |
| S2H13_7jv6 | 338 | 0.281 | 0.474 | 0.470 | -0.065 | -0.012 | 0.170 | 0.454 | 0.282 | 0.162 |
| S2H14_7jx3 | 476 | 0.433 | 0.291 | 0.430 | 0.221 | 0.041 | -0.040 | 0.208 | 0.097 | 0.039 |
| S2H97_7m7w | 322 | 0.370 | 0.254 | 0.450 | 0.151 | 0.204 | 0.040 | 0.278 | -0.060 | 0.029 |
| S2X259_7m7w | 430 | 0.332 | 0.180 | 0.360 | 0.016 | 0.014 | -0.020 | 0.045 | 0.050 | -0.004 |
| S2X35_7r6w | 394 | 0.435 | 0.293 | 0.460 | 0.027 | 0.039 | 0.060 | 0.279 | 0.020 | 0.047 |
| S304_7jx3 | 407 | 0.322 | 0.361 | 0.350 | -0.139 | 0.008 | 0.170 | 0.274 | -0.012 | 0.032 |
| S309_7r6w | 315 | 0.440 | 0.353 | 0.220 | 0.166 | 0.095 | 0.130 | 0.002 | -0.410 | -0.027 |
| VRC01_5ies | 679 | 0.517 | 0.071 | 0.230 | 0.078 | 0.208 | 0.140 | 0.088 | -0.166 | -0.057 |
| VRC34-01_5i8h | 513 | 0.274 | 0.105 | 0.190 | -0.159 | -0.004 | 0.140 | -0.054 | 0.092 | 0.023 |
| WIBP-2B11_7e5y | 278 | 0.221 | 0.275 | 0.250 | 0.064 | 0.187 | -0.070 | 0.162 | -0.019 | 0.244 |
| ZV-67_5kvg | 570 | 0.255 | 0.032 | 0.120 | -0.002 | -0.002 | 0.020 | -0.062 | -0.013 | 0.007 |
| m102-4_6cmg | 1016 | 0.008 | 0.056 | 0.160 | -0.079 | 0.005 | 0.040 | 0.040 | -0.019 | 0.000 |
| nAH1-3_7txz | 836 | 0.386 | 0.123 | 0.380 | 0.124 | 0.127 | 0.220 | 0.181 | 0.215 | -0.019 |
| tanezumab_4edw | 980 | 0.232 | -0.038 | 0.220 | 0.306 | 0.231 | -0.230 | -0.080 | -0.267 | 0.026 |
| MEAN | 568.343 | 0.283 | 0.197 | 0.253 | -0.024 | 0.033 | 0.008 | 0.112 | 0.013 | 0.035 |

Table 1: Spearman correlation coefficients between the prediction scores and the experimental DMS values for each DMS protein and each predictor.

| DMS DATA | # mut | ACC | BeAt MuSiC | FoldX | CI | LOR | mCSM AB | SAAMBE 3D | SaProt | Korpm |
|---|---|---|---|---|---|---|---|---|---|---|
| 1-18_6udj | 690 | 0.600 | 0.557 | 0.753 | 0.510 | 0.553 | 0.480 | 0.531 | 0.601 | 0.584 |
| 106E6_6n1w | 551 | 0.539 | 0.665 | 0.705 | 0.268 | 0.466 | 0.582 | 0.235 | 0.579 | 0.531 |
| 121F_7uov | 482 | 0.696 | 0.615 | 0.638 | 0.768 | 0.719 | 0.350 | 0.366 | 0.484 | 0.592 |
| 17D4_6n1v | 475 | 0.725 | 0.648 | 0.597 | 0.170 | 0.313 | 0.777 | 0.367 | 0.691 | 0.419 |
| 256A_6p95 | 1257 | 0.670 | 0.456 | 0.731 | 0.427 | 0.452 | 0.525 | 0.546 | 0.511 | 0.538 |
| 372D_7uot | 967 | 0.467 | 0.574 | 0.481 | 0.575 | 0.584 | 0.499 | 0.455 | 0.387 | 0.509 |
| 377H_5vk2 | 1346 | 0.781 | 0.524 | 0.796 | 0.394 | 0.416 | 0.385 | 0.755 | 0.532 | 0.581 |
| 3BN-1074_5t3z | 475 | 0.518 | 0.360 | 0.806 | 0.453 | 0.456 | 0.499 | 0.700 | 0.502 | 0.706 |
| 89F_7uot | 731 | 0.504 | 0.544 | 0.543 | 0.711 | 0.692 | 0.342 | 0.460 | 0.482 | 0.558 |
| AZD1061_7l7e | 395 | 0.873 | 0.606 | 0.733 | 0.529 | 0.593 | 0.440 | 0.768 | 0.638 | 0.783 |
| AZD8895_7l7d | 345 | 0.960 | 0.870 | 0.943 | 0.308 | 0.485 | 0.355 | 0.889 | 0.568 | 0.277 |
| BD55-5840_7wrz | 88 | 0.857 | 0.775 | 0.875 | 0.842 | 0.842 | 0.554 | 0.622 | 0.383 | 0.479 |
| C002_7k8s | 578 | 0.776 | 0.555 | 0.724 | 0.257 | 0.393 | 0.299 | 0.813 | 0.647 | 0.567 |
| C105_6xcm | 430 | 0.873 | 0.475 | 0.529 | 0.812 | 0.559 | 0.424 | 0.530 | 0.675 | 0.612 |
| C110_7k8v | 345 | 0.589 | 0.557 | 0.519 | 0.516 | 0.477 | 0.648 | 0.614 | 0.481 | 0.258 |
| C119_7k8w | 243 | 0.365 | 0.423 | 0.315 | 0.524 | 0.504 | 0.508 | 0.559 | 0.704 | 0.401 |
| C121_7k8x | 642 | 0.553 | 0.626 | 0.705 | 0.485 | 0.498 | 0.411 | 0.589 | 0.297 | 0.858 |
| C135_7k8z | 220 | 0.565 | 0.591 | 0.808 | 0.308 | 0.302 | 0.297 | 0.795 | 0.759 | 0.295 |
| C144_7k90 | 554 | 0.800 | 0.728 | 0.881 | 0.219 | 0.418 | 0.508 | 0.669 | 0.485 | 0.677 |
| C179_4hlz | 855 | 0.482 | 0.621 | 0.530 | 0.437 | 0.309 | 0.444 | 0.500 | 0.620 | 0.505 |
| COV2-2130_8d8q | 650 | 0.734 | 0.623 | 0.771 | 0.484 | 0.540 | 0.511 | 0.776 | 0.622 | 0.453 |
| COV2-2196_8d8q | 495 | 0.744 | 0.810 | 0.880 | 0.313 | 0.461 | 0.287 | 0.776 | 0.576 | 0.323 |
| COVA2-04_7jmo | 292 | 0.677 | 0.678 | 0.574 | 0.578 | 0.472 | 0.599 | 0.614 | 0.679 | 0.411 |
| CR3022_6zlr | 456 | 0.809 | 0.697 | 0.805 | 0.423 | 0.411 | 0.424 | 0.699 | 0.488 | 0.608 |
| Cetuximab_1yy9 | 659 | 0.621 | 0.555 | 0.572 | 0.374 | 0.464 | 0.529 | 0.625 | 0.568 | 0.441 |
| D441_1mlc | 627 | 0.522 | 0.594 | 0.578 | 0.436 | 0.523 | 0.531 | 0.554 | 0.433 | 0.661 |
| DF1W314_6mph | 722 | 0.663 | 0.670 | 0.725 | 0.258 | 0.359 | 0.493 | 0.355 | 0.682 | 0.603 |
| EDE1-C10_5h37 | 1501 | 0.497 | 0.468 | 0.464 | 0.391 | 0.396 | 0.556 | 0.459 | 0.468 | 0.642 |
| FAB-B_8ath | 399 | 0.521 | 0.692 | 0.637 | 0.445 | 0.415 | 0.591 | 0.502 | 0.546 | 0.344 |
| FI6V3-H3_3ztj | 513 | 0.550 | 0.834 | 0.648 | 0.629 | 0.571 | 0.634 | 0.757 | 0.434 | 0.297 |
| FI6V_3ztn | 703 | 0.476 | 0.466 | 0.521 | 0.384 | 0.307 | 0.442 | 0.435 | 0.624 | 0.500 |
| FP16-02_6cdi | 608 | 0.890 | 0.792 | 0.796 | 0.140 | 0.326 | 0.511 | 0.378 | 0.568 | 0.370 |
| FP20-01_6cde | 665 | 0.866 | 0.847 | 0.800 | 0.115 | 0.271 | 0.562 | 0.295 | 0.665 | 0.385 |
| G6_27_30A_4zff | 475 | 0.668 | 0.439 | 0.754 | 0.676 | 0.675 | 0.489 | 0.314 | 0.322 | 0.475 |
| G6_27_30A_4zfg | 471 | 0.568 | 0.536 | 0.716 | 0.373 | 0.571 | 0.273 | 0.690 | 0.337 | 0.559 |

| DMS DATA | # mut | ACC | BeAt MuSiC | FoldX | CI | LOR | mCSM AB | SAAMBE 3D | SaProt | Korpm |
|---|---|---|---|---|---|---|---|---|---|---|
| HENV26_6vy5 | 1144 | 0.573 | 0.412 | 0.792 | 0.371 | 0.380 | 0.371 | 0.516 | 0.491 | 0.471 |
| HENV32_6vy4 | 814 | 0.455 | 0.605 | 0.654 | 0.683 | 0.647 | 0.715 | 0.737 | 0.458 | 0.695 |
| LY-CoV016_7c01 | 510 | 0.767 | 0.612 | 0.779 | 0.763 | 0.580 | 0.416 | 0.636 | 0.639 | 0.612 |
| LY-CoV1404_7mmo | 170 | 0.480 | 0.716 | 0.667 | 0.313 | 0.390 | 0.514 | 0.488 | 0.408 | 0.312 |
| LY-CoV488_7kmh | 207 | 0.618 | 0.512 | 0.703 | 0.495 | 0.577 | 0.480 | 0.668 | 0.474 | 0.590 |
| LY-CoV555_7kmg | 431 | 0.901 | 0.557 | 0.846 | 0.142 | 0.261 | 0.475 | 0.726 | 0.468 | 0.416 |
| MZ4_6niu | 741 | 0.598 | 0.468 | 0.778 | 0.391 | 0.421 | 0.610 | 0.676 | 0.567 | 0.614 |
| OPV12_6ot1 | 532 | 0.813 | 0.631 | 0.764 | 0.273 | 0.430 | 0.633 | 0.273 | 0.586 | 0.474 |
| OPV20_6osy | 570 | 0.653 | 0.730 | 0.619 | 0.227 | 0.342 | 0.462 | 0.311 | 0.687 | 0.511 |
| PG9_3u4e | 437 | 0.721 | 0.532 | 0.480 | 0.538 | 0.547 | 0.484 | 0.704 | 0.500 | 0.534 |
| PGT121_7uoj | 399 | 0.525 | 0.549 | 0.795 | 0.613 | 0.561 | 0.647 | 0.711 | 0.666 | 0.575 |
| PGT145_5v8l | 646 | 0.542 | 0.699 | 0.800 | 0.469 | 0.423 | 0.494 | 0.585 | 0.736 | 0.345 |
| PGT151_5fuu | 817 | 0.628 | 0.414 | 0.739 | 0.450 | 0.456 | 0.542 | 0.349 | 0.442 | 0.478 |
| REGN10933_6xdg | 385 | 0.855 | 0.894 | 0.912 | 0.277 | 0.481 | 0.406 | 0.886 | 0.725 | 0.205 |
| REGN10987_6xdg | 265 | 0.766 | 0.823 | 0.771 | 0.209 | 0.355 | 0.564 | 0.782 | 0.563 | 0.426 |
| S139_4gms | 741 | 0.773 | 0.462 | 0.769 | 0.284 | 0.331 | 0.688 | 0.712 | 0.415 | 0.511 |
| S2D106_7r7n | 470 | 0.832 | 0.769 | 0.798 | 0.101 | 0.301 | 0.295 | 0.768 | 0.872 | 0.312 |
| S2E12_7r6x | 341 | 0.924 | 0.856 | 0.846 | 0.375 | 0.448 | 0.193 | 0.792 | 0.570 | 0.358 |
| S2H13_7jv6 | 338 | 0.842 | 0.884 | 0.886 | 0.365 | 0.533 | 0.696 | 0.688 | 0.726 | 0.252 |
| S2H14_7jx3 | 476 | 0.708 | 0.759 | 0.817 | 0.517 | 0.525 | 0.388 | 0.717 | 0.638 | 0.395 |
| S2H97_7m7w | 322 | 0.773 | 0.620 | 0.838 | 0.751 | 0.795 | 0.469 | 0.717 | 0.446 | 0.307 |
| S2X259_7m7w | 430 | 0.868 | 0.683 | 0.923 | 0.442 | 0.446 | 0.468 | 0.375 | 0.554 | 0.672 |
| S2X35_7r6w | 394 | 0.936 | 0.661 | 0.932 | 0.408 | 0.453 | 0.501 | 0.523 | 0.498 | 0.289 |
| S304_7jx3 | 407 | 0.912 | 0.781 | 0.850 | 0.395 | 0.388 | 0.670 | 0.713 | 0.517 | 0.525 |
| S309_7r6w | 315 | 0.900 | 0.853 | 0.957 | 0.724 | 0.734 | 0.433 | 0.561 | 0.184 | 0.569 |
| VRC01_5ies | 679 | 0.662 | 0.555 | 0.648 | 0.493 | 0.643 | 0.626 | 0.505 | 0.328 | 0.594 |
| VRC34-01_5i8h | 513 | 0.501 | 0.528 | 0.514 | 0.353 | 0.481 | 0.571 | 0.477 | 0.510 | 0.608 |
| WIBP-2B11_7e5y | 278 | 0.646 | 0.769 | 0.701 | 0.587 | 0.650 | 0.448 | 0.671 | 0.568 | 0.346 |
| ZV-67_5kvg | 570 | 0.623 | 0.413 | 0.586 | 0.445 | 0.289 | 0.491 | 0.189 | 0.562 | 0.342 |
| m102-4_6cmg | 1016 | 0.575 | 0.537 | 0.591 | 0.447 | 0.475 | 0.485 | 0.542 | 0.470 | 0.489 |
| nAH1-3_7txz | 836 | 0.854 | 0.544 | 0.869 | 0.587 | 0.551 | 0.745 | 0.707 | 0.681 | 0.539 |
| tanezumab_4edw | 980 | 0.557 | 0.499 | 0.628 | 0.643 | 0.537 | 0.359 | 0.518 | 0.406 | 0.573 |
| MEAN | 568.3 | 0.683 | 0.624 | 0.718 | 0.443 | 0.481 | 0.494 | 0.585 | 0.543 | 0.489 |

Table 2: ROC-AUC between the prediction scores and the binarized DMS values (destabilizing and neutral classes as defined in the main text) for each DMS protein and each predictor.

[13] Tiphanie Pruvost, Magali Mathieu, Steven Dubois, Bernard Maillère, Emmanuelle Vigne, and Hervé Nozach. Deciphering cross-species reactivity of lamp-1 antibodies using deep mutational epitope mapping and alphafold. In *MAbs*, volume 15, page 2175311. Taylor & Francis, 2023.

[14] Patrizia Ferraboschi, Samuele Ciceri, and Paride Grisenti. Applications of Lysozyme, an Innate Immune Defense Factor, as an Alternative Antibiotic. *Antibiotics*, 10(12):1534, December 2021.

[15] Charles M. Forsyth, Veronica Juan, Yoshiko Akamatsu, Robert B. DuBridge, Minhtam Doan, Alexander V. Ivanov, Zhiyuan Ma, Dixie Polakoff, Jennifer Razo, Keith Wilson, and David B. Powers. Deep mutational scanning of an antibody against epidermal growth factor receptor using mammalian cell display and massively parallel pyrosequencing. *mAbs*, 5(4):523–532, 2013. PMID: 23765106.

[16] Patrick Koenig, Chingwei V Lee, Sarah Sanowar, Ping Wu, Jeremy Stinson, Seth F Harris, and Germaine Fuh. Deep sequencing-guided design of a high affinity dual specificity antibody to target two angiogenic factors in neovascular age-related macular degeneration. *Journal of Biological Chemistry*, 290(36):21773–21786, 2015.
